# Activation of perivascular thermogenic fat improves aortic stenosis

**DOI:** 10.64898/2026.10.13.758767

**Authors:** Staffan Hildebrand, Klaus Wunderling, Zuzanna Siwik-Róg, Alexander Schmidt, Jorge A. Barrero, Nicolás Gómez-Banoy, Laia Reverte-Salisa, Benjamin Freyter, Sebastian Zimmer, Georg Nickenig, Paul Cohen, Alexander Pfeifer

## Abstract

**Aim:** Aortic stenosis (AS) is a common condition with exponential increases in incidence in aging populations. Perivascular adipose tissue (PVAT) forms the outer layer of the aortic wall and has – especially in the thoracic area – characteristics of thermogenic adipose tissue, which dissipates energy and releases endocrine factors. Here, we set out to investigate the role of thermogenic PVAT in AS.

**Methods and results:** We studied the role of (PVAT) in the development of AS in a murine model of AS. To specifically delineate the role of PVAT activity in AS, we used a pharmacological β3-adrenergic agonist as well as cold exposure in combination with a genetic mouse model with deficient PVAT development. To investigate the role of thermogenic adipose tissue in human AS, we used ^18^F-FDG PET-CT imaging. We found that pharmacological activation of thermogenic adipose tissue resulted in an improvement of AS, but not in mice genetically deficient in PVAT. Similarly, PVAT was found to be protective against AS during physiological stimulation of PVAT using cold exposure. Mechanistically, PVAT activation resulted in enhanced aortic distensibility and cellular remodeling of the aortic wall, possibly via release of polyunsaturated fatty acids and oxylipins from perivascular adipocytes. Finally, analysis of thermogenic fat activity in humans revealed that mediastinal thermogenic fat activity correlated negatively with diagnostic criteria of AS severity, suggesting an important role for PVAT activity in human AS.

**Conclusion:** Our data indicate that PVAT activation reduces aortic stenosis and supports aortic remodeling to improve ascending aorta hemodynamics. PVAT activation might be a novel therapeutic target to ameliorate AS progression.

**Translational perspective:** Currently, there are no effective strategies for treating AS other than valve replacement. Since the molecular mechanisms underlying the disease progression remain unclear, there are no pharmacological agents approved for either treatment or prevention of this disease. Furthermore, AS development cannot currently be predicted and is often only diagnosed when patients develop symptoms, which may be 10-20 years after disease onset. Therefore, identifying novel mechanisms that improve aortic valve function and prevent AS is of high medical need. This study identifies activation of PVAT-dependent aortic remodeling as a potential new therapeutic approach to improve or prevent AS.

## Introduction

Obesity is a major risk factor for cardiovascular diseases ^1^ including myocardial infarction and stroke. Although obesity is defined as an abnormal increase in body fat, the role of different types of fat in cardiovascular disease varies. White adipose tissue (WAT) is the largest store of energy in the human body and its expansion during obesity is a major cause of so-called metaflammation, a state of chronic inflammation that affects other organs ^2, 3^. In contrast, brown adipose tissue (BAT) has the ability to dissipate energy as heat (thermogenesis) by mitochondrial uncoupling. This is achieved by Uncoupling protein 1 (UCP1), which uncouples ATP production from the respiratory chain by disrupting the proton gradient across the inner mitochondrial membrane ^4^. Additionally, several other futile cycling mechanisms have recently been identified that can drive thermogenesis even in the absence of UCP1 ^5^. In addition to their role in thermogenesis, brown adipocytes secrete numerous active substances that have been demonstrated to have both para- and endocrine roles ^6^. Importantly, thermogenic fat activity correlates with lower odds ratio for cardiovascular disease (including coronary artery disease and hypertension) and improved metabolic health^7^.

Aortic stenosis (AS) is a degenerative disease with a complex and understudied etiology, and is characterized by narrowing and calcification of the aortic valve. If untreated, AS can lead to heart failure, and the 2-year mortality rate after onset of symptoms is as high as 50% ^8^. Currently, there is no effective pharmacological therapy, and aortic valve replacement remains the only effective therapeutic approach. Perivascular adipose tissue (PVAT) is directly adjacent to the aortic wall and is an integral part of the aortic organ. PVAT around the thoracic aorta is a type of thermogenic adipose tissue that expresses high levels of UCP1, and has a strong transcriptomic, proteomic, and metabolic similarity to classical BAT ^9–11^. Like classical BAT, PVAT secretes several paracrine factors, some of which have vasoactive properties and can modulate different aspects of aortic function ^12^. So far, however, little is known about how PVAT activation affects AS development.

Here, we studied the interaction between PVAT and AS using a genetic mouse model with defective PVAT development in conjunction with a wire-injury model of AS. Activation of PVAT improves valvular function in AS as reflected by diminished peak velocities and transvalvular pressure gradients, as well as reduced histological valve stenosis morphology. Mechanistically, we found that prolonged PVAT activation leads to aortic remodeling that increases aortic compliance and improves AS. Finally, human mediastinal thermogenic fat activity correlated significantly with both aortic valve area and peak transvalvular blood flow velocity, indicating that PVAT activity is linked to human aortic valve function.

## Methods

### Human data

Patients from Memorial Sloan Kettering Cancer Center (MSKCC) with detectable thermogenic fat in their ^18^F-FDG-PET CT scan and measurements of metabolic activity were screened for the diagnosis of aortic stenosis (AS) (ICD code I35.0). Eleven patients with AS diagnosis were identified. Metabolic activity in the mediastinal thermogenic fat depot was quantified utilizing the TriDFusion method ^13^. The aortic valve area (cm^2^) and maximum velocity of aortic valve flow (cm/s) were extracted from the echocardiogram reports of these patients. The correlation between these echocardiographic parameters and mediastinal BAT metabolic activity was assessed after logarithmic transformation for data normalization using a Pearson correlation test. The study followed institutional guidelines and was approved by the Institutional Review Boards of The Rockefeller University and MSKCC and conformed to the principles outlined in the Declaration of Helsinki. Due to the retrospective nature of this study, the requirement for informed consent was waived.

### Animals

Sm22α^Cre/+^ (B6.Cg-Tg(Tagln-Cre)1Her/J) mice were kindly provided by Robert Feil and Joachim Herz ^14^ and crossed with PPARγ^fl/fl^ (B6.129-Pparg^tm2Rev^/J) mice, kindly provided by Irmgard Förster, to generate PVAT-deficient mice. All mice were maintained on a daily cycle of 12 h light (06:00–18:00) and 12 h darkness (18:00– 06:00), at ambient room temperature (23 ± 1 °C) and 40–50% humidity if not otherwise stated, and were allowed free access to chow and water. Wild-type C57BL/6J mice were purchased from Charles River. Mice were euthanized by cervical dislocation or by exsanguination following anesthesia via intraperitoneal injection of ketamine (150 mg/kg) and xylazine (16 mg/kg). For wire injury surgeries, mice were anesthetized by intraperitoneal injection of ketamine (150 mg/kg) and xylazine (16 mg/kg), and a single intraperitoneal administration of buprenorphine (0.1 mg/kg) followed by buprenorphine supplementation in the drinking water (10 µg/ml) was used as analgesia. For osmotic minipump implantation, mice were anesthetized by isoflurane inhalation (3-5% initiation, 1-2% maintenance), and a single intraperitoneal administration of buprenorphine (0.1 mg/kg) followed by supplementation of metamizole in the drinking water (1.33 mg/ml) was used as analgesia. The use of animals conformed to the guidelines from Directive 2010/63/EU of the European Parliament on the protection of animals used for scientific purposes. All animal experiments were approved by Landesamt für Verbraucherschutz und Ernährung, Nordrhein-Westfalen, Germany, under the governmental approval numbers 84-02.04.2016.A202 and M2026-318.

## Results

### Pharmacological activation of PVAT ameliorates AS

To determine the relationship between PVAT activity and AS, we investigated the functional effect of PVAT activation on AS development in the graded wire injury model of murine AS. To distinguish between PVAT-mediated effects and non-PVAT-mediated effects, we performed these experiments in mice lacking PVAT and control mice. For this, we used the previously described genetic model of PVAT ablation (PPARγ^fl/fl^ Sm22a^Cre/+^, PVAT-) ^10^. These mice display strongly reduced PVAT mass compared to control mice (PPARγ^fl/fl^ Sm22a^+/+^, PVAT+), while retaining fully functional classical interscapular BAT (Figure 1A, Supplemental Figure 1A-B). To analyze the effect of PVAT activation on AS, we treated these mice with the β3-specific adrenergic agonist CL316,243 (CL, 1 mg/kg), which specifically activates thermogenic adipose tissues (Figure 1B), for the entire duration of stenosis development after wire injury. Immunohistological analysis showed that CL treatment increased UCP1 expression in both PVAT and inguinal WAT, indicating robust activation of the thermogenic program (i.e. “browning”) in PVAT (Supplemental Figure 1B-C). Doppler echocardiography-based assessment of aortic stenosis showed that PVAT activation by CL treatment led to both significantly reduced peak aortic jet velocity and peak transvalvular pressure gradient four weeks after wire injury in PVAT+ mice, strongly suggesting that activation of thermogenic fat ameliorates AS development (Figure 1C). Strikingly, CL did not reduce peak velocity and peak gradient in PVAT- mice (Figure 1D). Further detailed histological analysis revealed that PVAT+ mice treated with CL had significantly reduced leaflet area as determined by H&E staining (Figure 1E,G). Additionally, picrosirius red staining of the valve leaflets revealed significantly reduced fibrosis, while immunostaining with the pan-macrophage marker CD68 showed no difference in leaflet macrophage infiltration. (Figure 1E,G). Conversely, CL treatment resulted in significantly increased leaflet area as well as increased picrosirius red birefringence in PVAT- mice. CD68 staining did not show any changes in valvular macrophage infiltration after CL treatment. (Figure 1F,H). Together, these data strongly suggest that thermogenic activation of PVAT is protective during AS development and reduces AS severity.

**Figure 1:**
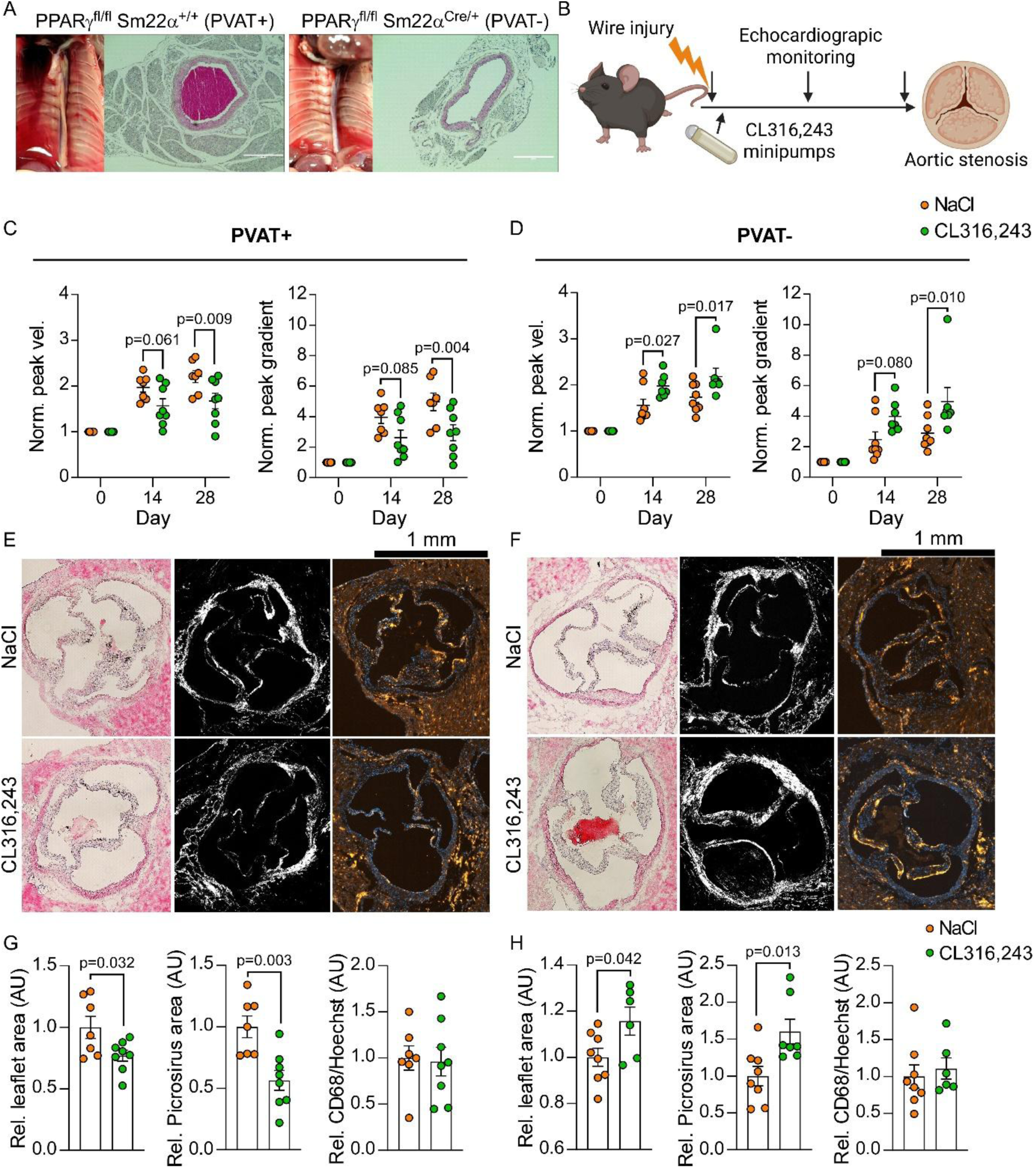
Treatment with CL316,243 reduces AS in a PVAT-dependent manner. **A:** Macroscopic photographs and H&E stainings showing loss of PVAT in PPARγ^fl/fl^ Sm22a^Cre/+^ (PVAT-) mice. **B:** Schematic depicting experimental setup. **C:** Echocardiographic analysis of valve hemodynamics in PVAT+ mice at different timepoints. n=7-8. Two-way ANOVA and Sidak’s post-hoc test. **D:** Echocardiographic analysis of valve hemodynamics in PVAT- mice at different timepoints. n=7-8. Two-way ANOVA. **E:** H&E, Picrosirius red, and CD68-staining in aortic valves from PVAT+ mice following wire injury and treatment with either CL or NaCl. Scale bar: 1 mm. **F:** H&E, Picrosirius red, and CD68-staining in aortic valves from PVAT- mice following wire injury and treatment with either CL or NaCl. Scale bar: 1 mm. **G:** Quantification of leaflet area, picrosirius red birefringence, and CD68/Hoechst area in aortic leaflets of PVAT+ mice. n=7-8. t-test. **H:** Quantification of leaflet area, picrosirius red birefringence, and CD68/Hoechst area in aortic leaflets of PVAT- mice. n=6-8. t-test. **Alt text:** Graphs showing how activation of thermogenic fat improves aortic stenosis development in mice in a PVAT-dependent manner.

### Physiological PVAT activation protects against AS

Next, we used cold exposure as a physiological stimulus for thermogenic fat activation and studied its effect on AS development. Exposure to cold is known to cause peripheral vasoconstriction and increased central pulse pressure ^15^, which is linked to increased mechanical stress on the aortic valve leaflets ^16^. Additionally, activation of thermogenic fat increases oxygen demand, leading to increased cardiac output. Since we saw a protective effect of pharmacological PVAT activation on AS, we tested if PVAT protects the aortic valves during cold stress. Therefore, we subjected PVAT- and PVAT+ mice to wire injury-induced AS with concomitant exposure to cold for the entire duration of AS development (Supplemental Figure 2A). Cold-exposed PVAT- mice showed a significant worsening of AS development, with increased peak velocity, pressure gradient, and aortic valve area (Supplemental Figure 2B-C). Additionally, immunohistological staining for CD68 showed that PVAT- mice had increased macrophage infiltration in the aortic leaflets, suggesting increased inflammation (Supplemental Figure 2D). Both PVAT+ and PVAT- mice exhibited increased heart mass following the five weeks of housing at 8°C (Supplemental Figure 2E). In contrast to PVAT- mice, echocardiographic analyses revealed that cold-exposure did not significantly affect either peak jet velocity or peak transvalvular pressure gradient after wire injury in mice with intact PVAT (Supplemental Figure 2F). Histological assessment of aortic leaflet area also did not show any differences between cold-exposed and control PVAT+ mice. (Supplemental Figure 2G). In summary, these data show that cold stress significantly worsens the progression of AS in the absence of functional PVAT, suggesting that PVAT protects the aortic valves during cold stress.

### Effects of PVAT activation on aortic hemodynamics

To investigate the mechanism behind the protective effects of PVAT activation on AS, we focused on the effect of PVAT on aortic hemodynamics. We found a significant increase of aortic diameter change during the cardiac cycle after activation of thermogenic fat with CL (Figure 2A-B). Interestingly, this increase was only observed in mice with PVAT and not in PVAT- mice (Figure 2B). Stroke volume and cardiac output were not significantly different between CL-treated PVAT+ and PVAT- mice (Figure 2C), suggesting that PVAT activation enhances compliance of the aorta and thereby the Windkessel function, thereby potentially creating a more favorable hemodynamic environment for the aortic valves. Computational as well as *ex vivo* studies have shown that increased stiffness of the aorta and hypertension leads to heightened strain on the aortic valves during the cardiac cycle ^17–19^.

**Figure 2:**
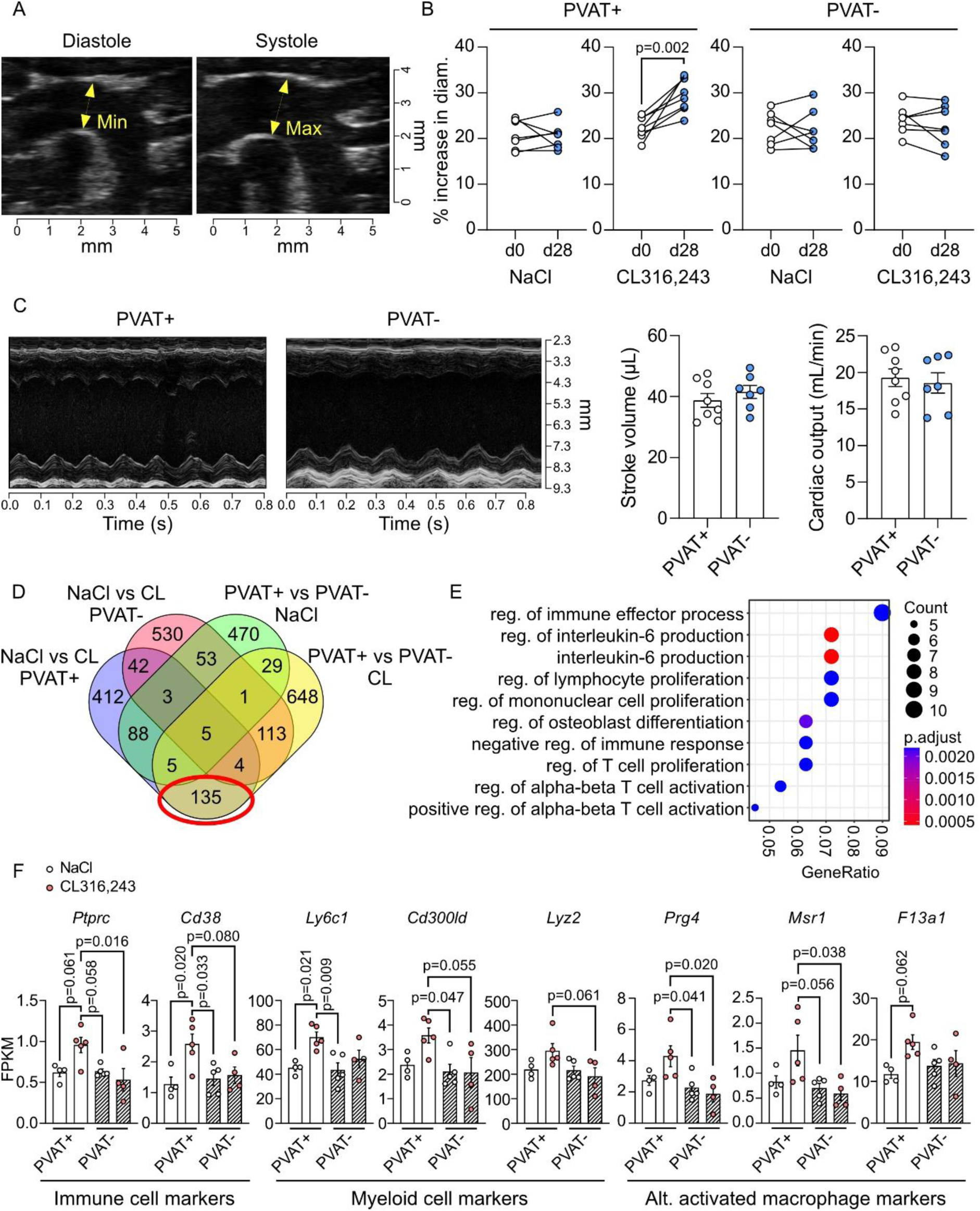
CL treatment increases aortic distensibility and causes increased expression of immune cells markers in the ascending aorta. **A:** Assessment of aortic strain in PVAT+ and PVAT- mice with AS. B: Changes in aortic strain during CL treatment in PVAT+ and PVAT- mice with AS. n=7-8. t-test. C: Stroke volume and cardiac output in CL-treated PVAT+ and PVAT- mice. n=7-8. **D:** Venn diagram showing identification of DEGs regulated by CL in a PVAT-dependent manner (circled). **E:** GO analysis showing enrichment of immune cell regulation in the ascending aorta of CL-treated PVAT+ mice. **F:** Expression of pan-immune cell markers, myeloid cell markers, and markers of alternatively activated macrophages. n=4-5. One-way ANOVA with Tukey’s post-hoc test. **Alt text:** Graphs showing that PVAT activation results in increased aortic distensibility, which is associated with increased expression of markers for remodeling macrophages.

To further investigate what causes the effect of PVAT activation on aortic distensibility, we performed bulk mRNA sequencing of the ascending aorta from wire-injured PVAT+ and PVAT- mice with or without treatment with CL. To identify PVAT-dependent signals induced by CL, we filtered out differentially expressed genes (DEGs) that showed differences in expression between CL vs NaCl (vehicle) in PVAT+ mice, and between CL-treated PVAT+ and PVAT- mice (Figure 2D). Surprisingly, gene ontology analysis of the resulting hits showed increased signs of immune cell involvement (Figure 2E), and analysis of specific DEGs revealed an increased expression of the immune cell markers CD45 (*Ptprc*) and CD38 (*Cd38*) in the aortas from CL-treated PVAT+ mice (Figure 2F). Looking at markers specific for different immune cell types, we found increased expression of myeloid cell (*Ly6c1*, *Cd300ld*, and *Lyz2*) and alternatively activated/scavenger macrophage markers (*Prg4*, *F13a1*, and *Msr1*), suggesting an influx of these cell types in a PVAT-dependent manner during CL treatment (Figure 2F). Aortic remodeling macrophages are heavily involved in the regulation of aortic stiffness by regulating collagen expression by smooth muscle cells (SMC), and are characterized by expression of the hyaluronan receptor LYVE-1 ^20^. To further investigate alterations in aortic cell populations after PVAT activation, we performed single-nucleus sequencing (snSeq) of the aortas, including PVAT, from mice injected with CL or NaCl daily for 10 days. Since our bulk mRNA sequencing data suggested an increased accumulation of differentiated remodeling macrophages, we focused on this earlier time-point in order to also capture the influx of recruited monocytes. CL treatment led to extensive cellular remodeling of the aortic cell populations (Figure 3A-B). Aortas from CL-treated mice showed an increase in the proportion of committed brown preadipocytes and a reduction of white adipocytes, indicating robust activation of the PVAT (Figure 3B). Interestingly, CL treatment also led to a reduced population of synthetic SMC expressing fibrillar collagens, and an increase in myeloid cells (Figure 3B). Further analysis of the SMC compartment revealed reduced expression of the fibrillar collagen genes *Col1a1* and *Col3a1*, encoding type I and III collagens, which are critical determinants of aortic stiffness (Figure 3C-D). Detailed analysis of the immune cell compartment additionally revealed an increase in *Lyve1*^high^ macrophages as well as a strong increase in infiltrating monocytes, corroborating our bulk mRNA sequencing data (Figure 3E, Supplemental Figure 3A). To further validate the influx of macrophages, we injected mice with CL daily for 10 days and analyzed macrophage content in the ascending aorta by immunostaining. We found a significant increase in staining intensity for the macrophage marker CD68 in the adventitia of CL-injected mice (Figure 3F). Additionally, gene ontology analysis in genes regulated by CL in PVAT+ vs. PVAT- mice showed significant differences in pathways involving collagen distribution and extracellular matrix organization (Supplemental Figure 3B), further suggesting a PVAT-dependent remodeling of the aortic wall in response to CL treatment. Taken together, these results show that PVAT activation increases aortic distensibility which is associated with regulation of collagen expression in SMC and an influx of remodeling macrophages to the aortic wall.

**Figure 3:**
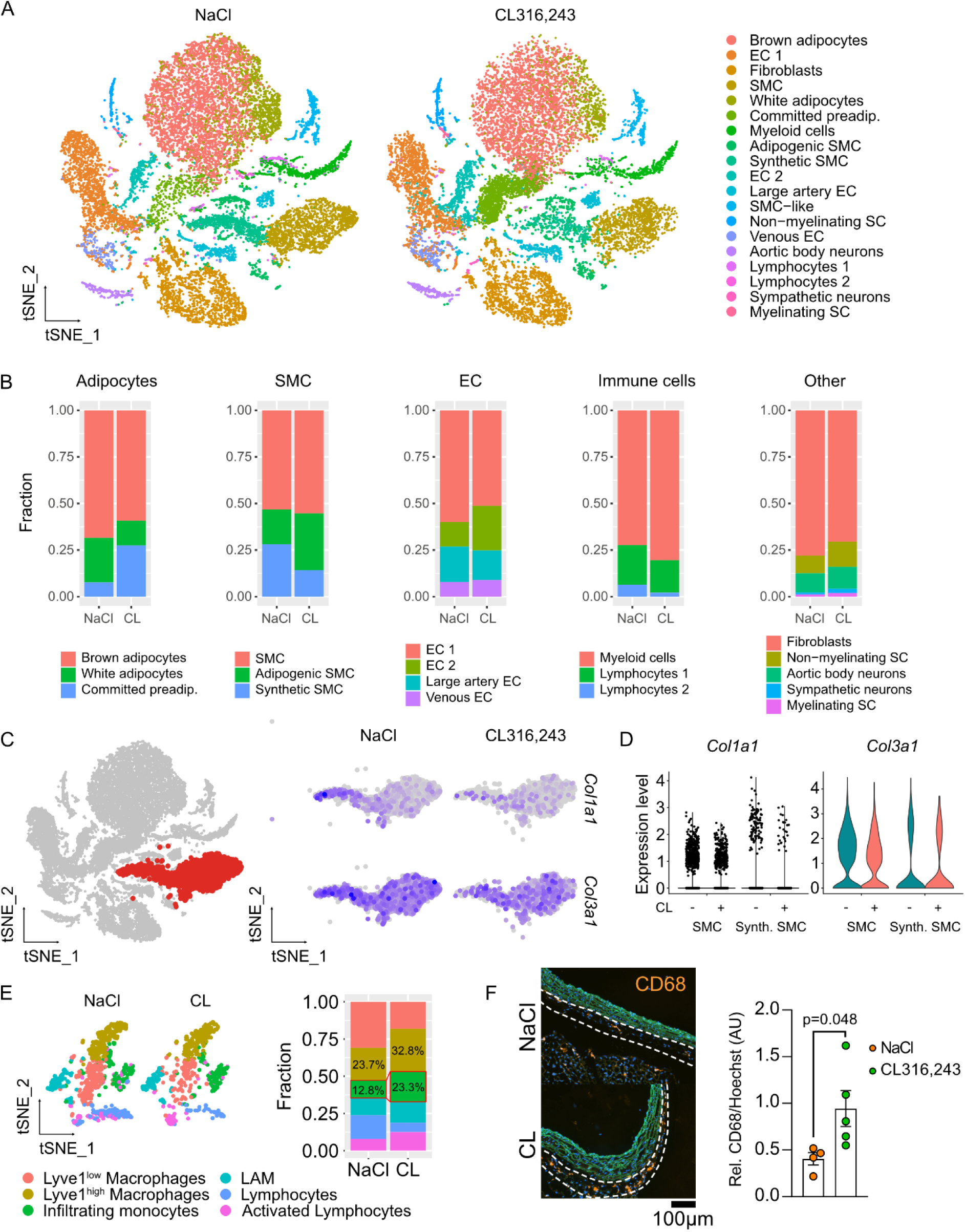
PVAT activation leads to cellular remodeling of the aorta. **A:** Dimensional reduction plots of snRNAseq data showing cellular populations in the aortas (including PVAT) of CL316,243- or saline-injected mice. **B:** Relative changes in the aortic cellular compartments following PVAT activation. **C:** Feature plots showing Col1a1 and Col3a1 in aortic SMC following CL316,243 treatment. **D:** Violin plots showing Col1a1 and Col3a1 in different SMC populations with or without Cl316,243 treatment. **E:** Dimensional reduction plots showing the different immune cell populations in aortas of NaCl- or CL316,243-injected mice. **F:** Immunostaining for CD68 in the ascending aortas of CL-injected or control mice. Orange: CD68, blue: Hoechst, green: autofluorescence. Scale bar: 100 µm. n=4-5. t-test. **Alt text:** Figures depicting changes in different cellular populations of mouse aortas following activation of thermogenic fat.

### Activated perivascular adipocytes secrete oxylipins and their precursors

Next, we aimed to identify the PVAT-derived mediator of macrophage influx in the aorta during thermogenic activation. As our mRNA sequencing data from the ascending aortas of CL-treated PVAT+ and PVAT- mice did not reveal any differential expression of chemokines upon CL treatment in PVAT+ mice (Supplemental Figure 4A), and since a major function of PVAT is lipid storage, we focused on lipid-derived factors. Interestingly, GSEA analysis of the mRNA transcriptome of PVAT from cold exposed (4°C) or control mice housed at 23°C revealed enrichment of pathways involved in oxylipin and eicosanoid signaling (Figure 4A). Furthermore, unsupervised clustering of arachidonic acid (AA) synthases showed a clear dysregulation of AA synthase expression in the PVAT of cold-exposed mice (Figure 4B). To study the oxylipin release from perivascular adipocytes, we isolated the stromal vascular fraction of thoracic PVAT and purified PDGFRα+ CD200-adipocyte precursors by tandem MACS. After adipogenic differentiation, the mature perivascular adipocytes (PVA) were stimulated with CL, and the supernatant was collected. Analysis of oxylipin species and polyunsaturated fatty acid (PUFA) precursors from CL- and vehicle-treated PVAs revealed significantly increased release of AA, docosahexaenoic acid (DHA), PGE2, 15-HETE, and 15-HETrE (Figure 4C-D). Strikingly, the most abundantly released lipids were the n-3 and n-6 PUFAs AA and DHA, which are heavily involved in both chemoattraction and macrophage polarization via their various enzymatic oxidation products. Whereas AA can be converted by arachidonate 5-lipoxygenase (5-LOX, encoded by *Alox5*) into the potent eicosanoid chemoattractants 5-HETE, 5-oxo-ETE, and LTB4 (the latter via leukotriene-A4 hydrolase), DHA can likewise be converted by 5-LOX to potent anti-inflammatory mediators such as Resolvins, Electrophilic oxo-derivatives (EFOX), and the newly identified Vascular-Leukocyte Resolvin (VL-Rv) ^21^. Analysis of single-cell sequencing data from murine aortas showed substantial expression of *Alox5* in immune cell clusters (Figure 4E), particularly in *Lyve1*-expressing macrophages (Figure 4F). We therefore explored the possibility of transcellular oxylipin biosynthesis in resident macrophages. Previous reports have shown that dormant 5-LOX in naïve bone-marrow derived macrophages (BMDM) can be activated by exogenous AA, and that these cells can synthesize 5-LOX products upon simultaneous treatment with AA and a calcium ionophore ^22^. We therefore tested if AA and/or DHA alone is sufficient to induce 5-LOX activity in alternatively activated BMDM, which phenotypically resemble aortic *Lyve1*^+^ macrophages ^20^. Interestingly, treatment with 40µM AA resulted in a rapid release of both 5-HETE and LTB_4_ in the supernatant. In contrast, neither metabolite was detectable in the supernatant of control BMDM (Figure 4G). Based on these data, we performed an untargeted analysis of lipid mediators from BMDM treated with AA, DHA, or both. Measuring supernatants of DHA-treated BMDM, we identified peaks corresponding to the 5-LOX-derived D-series Resolvins as well as 4-hydroxy-docosahexaenoic acid (4-HDHA), a 5-LOX-derived precursor for VL-Rv (Figure 4H, Supplemental Figure 4B). Treatment with AA, but not DHA, resulted in the production of the chemoattractant 5-HETE as well as the AA-derived pro-resolving mediator Lipoxin B4 (Figure 4H, Supplemental Figure 4B). Importantly, co-incubation of BMDM with AA and DHA resulted in the simultaneous production of 5-LOX-derived chemoattractants and pro-resolving mediators (Supplemental Figure 4B). These data suggest that PUFAs released from activated PVA is sufficient to induce simultaneous production of eicosanoid chemoattractant and pro-resolving lipid mediators in macrophages, which could explain the observed influx of remodeling macrophage populations in the aortic wall after PVAT activation (Figure 4I).

**Figure 4:**
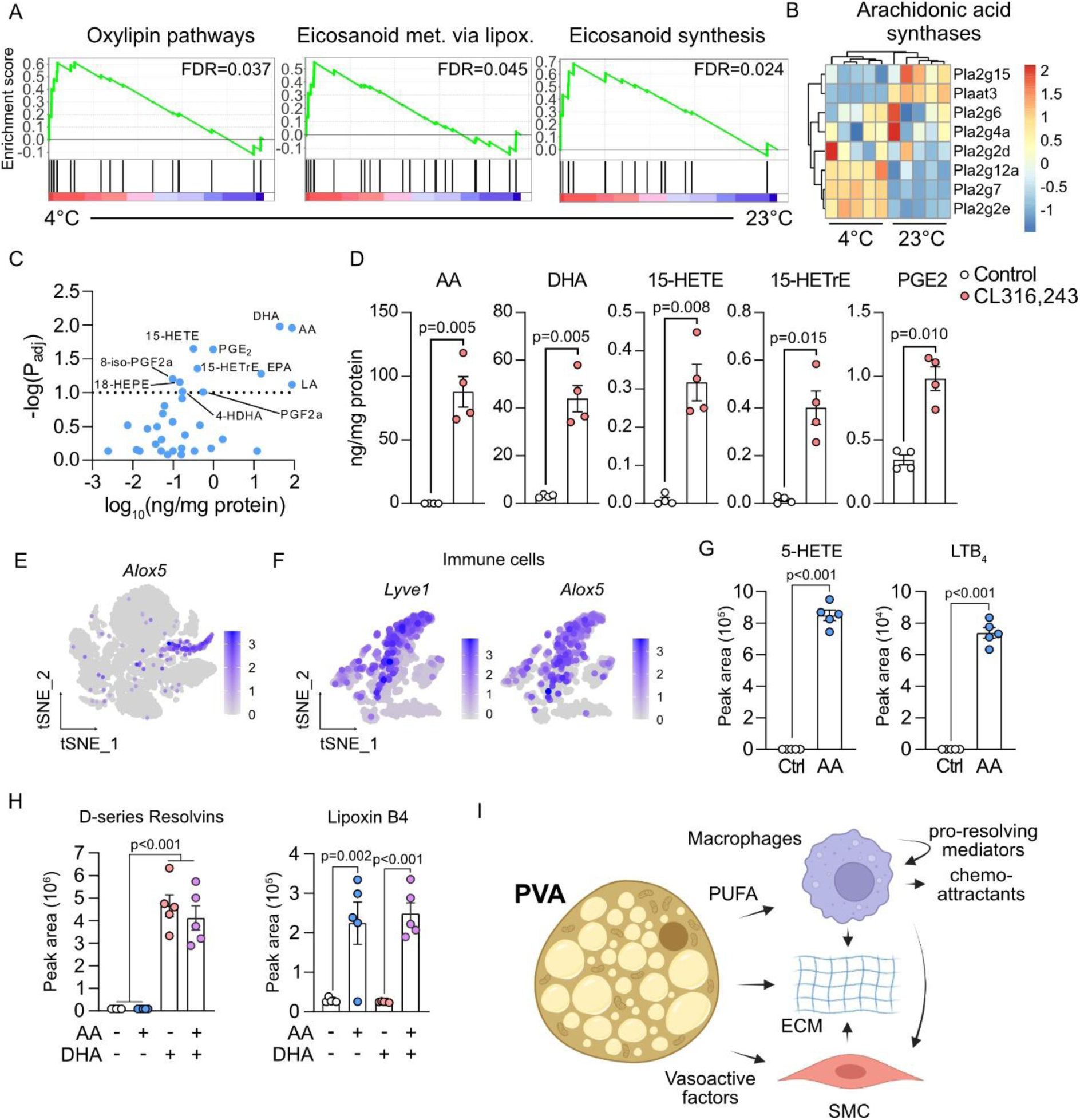
Activated PVA release PUFAs and oxylipins. **A:** GSEA analysis of wikipathways enrichment of the mRNA transcriptome of PVAT from cold-exposed or control mice. **B:** Unsupervised clustering of AA synthase expression in PVAT of cold-exposed or control mice. n=5. **C:** Scatter plot showing released oxylipins in the supernatant of CL-treated PVA and their abundance. n=4. **D:** Supernatant abundance of significantly increased oxylipin species after CL treatement of PVA. n=5. t-test. **E:** Feature plot showing Alox5 expression in aortic immune cells. **F:** Feature plots showing the expression of Alox5 in Lyve1^+^ cells of the aortic wall. **G:** Measurement of 5-HETE and LTB_4_ release in control- and AA-treated alternatively activated bone marrow-derived macrophages. n=5. t-test. **H:** Measurement of the 5-LOX products Resolvin D1-4 and Lipoxin B4 in alternatively activated BMDM treated with either AA or DHA or both. n=5. One-way ANOVA with Tukey’s post-hoc test. **I:** Schemed depicting working model of PVAT-mediated regulation of aortic remodeling. **Alt text:** Graphs showing involvement of perivascular fat-derived polyunsaturated fatty acids in the regulation of lipid mediators in macrophages.

### Mediastinal thermogenic fat activity is associated with improved echocardiographic parameters in AS patients

To investigate the role of thermogenic PVAT in AS in humans, we analyzed a large cohort of patients with thermogenic adipose tissue detectable by ^18^F-FDG PET-CT scans, which is the gold standard for assessing metabolic activity in thermogenic fat of humans. Patients diagnosed with mild to severe AS were identified (ICD code I35.0, Figure 5A), and mediastinal thermogenic fat metabolic activity (SUVmean*mL) – as a surrogate parameter for PVAT function – was correlated with AS severity, as measured by echocardiography. Mediastinal fat is adjacent to the ascending aorta and aortic valves and is the BAT depot that most closely mirrors murine aortic PVAT. Interestingly, we found a significant positive correlation between mediastinal thermogenic fat activity and aortic valve area (AVA), suggesting that high mediastinal thermogenic fat activity is associated with less severe stenosis in AS patients (Figure 5B). Additionally, peak transvalvular blood flow velocity showed a significant negative correlation with mediastinal thermogenic fat activity, further corroborating this notion (Figure 5C).

**Figure 5:**
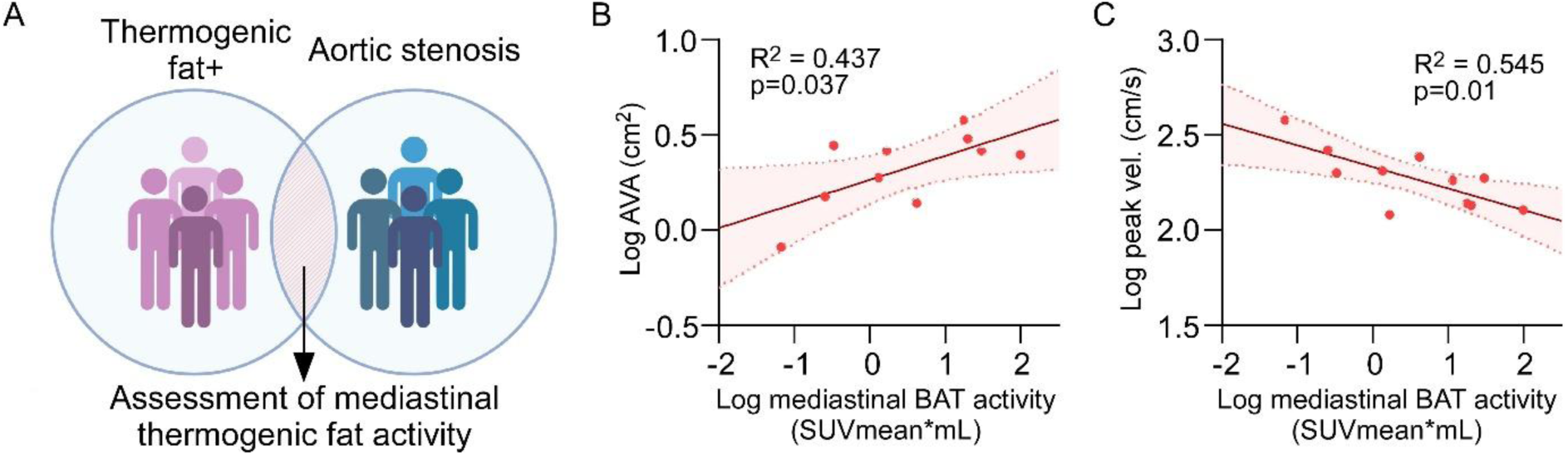
Mediastinal thermogenic fat activity is associated with reduced AS severity in humans. **A:** Scheme depicting the cohorts used in this study. Patients (n=8907) with measurements of thermogenic fat activity in ^18^F-FDG PET-CT scans were analyzed for AS. **B:** Correlation between aortic valve area (AVA) and mediastinal thermogenic fat metabolic activity in mediastinal fat. Simple linear regression. **C:** Correlation between transvalvular blood flow peak velocity and mediastinal thermogenic fat metabolic activity. Simple linear regression. **Alt text:** Figures depicting correlation of the activity of thermogenic fat in the chest cavity and improved hemodynamic parameters in patients with aortic stenosis.

## Discussion

The activity of thermogenic adipose tissues like PVAT has a well-established correlation with cardiometabolic health ^7^, but whether PVAT affects AS has so far not been studied. In this study, we investigated the role of PVAT activation in AS using a combination of surgical and genetic mouse models, as well as pharmacological and environmental challenges. We found that activation of thermogenic adipose tissue ameliorates AS in a PVAT-dependent manner, and that this is associated with an improved Windkessel effect in the ascending aorta.

The pressure-buffering properties of the aorta are critically important to regulate systolic blood pressure and pulse pressure, which are major factors for AS development ^23^, and decreased circumferential strain of the ascending aorta is associated with AS ^24^. Both *ex vivo* and *in silico* analyses have shown increased hemodynamic and mechanical stress on the valve leaflets under hypertensive conditions, supporting the notion of a mechano-etiology in AS ^19 18^. Our data complement these findings and indicate that improving aortic distensibility by PVAT activation reduces AS progression.

Thoracic aortic PVAT has long been recognized as a thermogenic fat BAT, displaying multilocular lipid droplets and high expression of UCP1, resembling classical BAT ^25^. The major physiological role of thermogenic adipose tissue is to maintain body temperature during cold stress ^26^. However, recent studies have shown that PVAT is largely dispensable for cold tolerance even in mice, suggesting a different principal role for PVAT ^10^. While PVAT also has an important function in regulating vascular tone ^27^, it has not been clear what role PVAT plays in large vessels and during AS development. Our findings outline a previously unrecognized role of PVAT as a depot for lipid mediators that support aortic remodeling to improve aortic distensibility under hemodynamic load. Adaptive remodeling of the aorta occurs in response to hemodynamic insults, and has primarily been studied in the context of pathogenesis, e.g. in hypertension and aortic aneurysm ^28, 29^. However, adaptive remodeling also occurs in the aorta under non-pathological circumstances that alter hemodynamic conditions, e.g. during prolonged exercise training, which leads to reduced aortic stiffness and central blood pressure ^30^.

The role of macrophages in this remodeling process is complex and highly dependent on pathogenesis as well as macrophage polarization state. For example, alternatively activated macrophages have repeatedly been shown to play a protective role in aortic aneurysm ^31–33^, while infiltration of inflammatory macrophages is a key driver for aortic aneurysm pathogenesis ^33–35^. The role of macrophages in aortic remodeling during hypertension has so far mostly been studied in artificially induced hypertension caused by Angiotensin II-infusion. This led to an initial influx of pro-inflammatory macrophages, followed by an accumulation of pro-remodeling macrophages at later stages ^36, 37^. In this context, macrophage infiltration was associated with increased fibrosis and loss of elastin ^37^. On the other hand, resident adventitial LYVE-1^+^ macrophages have recently been found to be essential in regulating aortic collagen production by SMC, thereby affecting both aortic collagen deposition and stiffness ^20^. These macrophages exhibit hallmark signs of the alternatively activated phenotype such as expression of CD163 and CD206 ^20^.Interestingly, we found evidence for a PVAT-dependent increase in abundance of alternatively activated macrophages in the aortas of mice treated with CL, which was associated with increased aortic distensibility and collagen remodeling. Single nucleus sequencing of aortas and PVAT also revealed an increased population of infiltrating monocytes as well as *Lyve1*^+^ macrophages at an earlier timepoint during CL treatment, indicating that PVAT activation leads to recruitment of remodeling LYVE-1^+^ macrophages in the aortic wall. Consistent with findings that LYVE-1^+^ macrophages regulate aortic stiffening by regulating collagen synthesis in SMC, we found a reduction in *Col1a1* and *Col3a1* production in aortic SMC following CL treatment. Mechanistically, we found that activated PVAs secrete large amounts of vasoactive PUFAs and oxylipins, including the n-6 fatty acid AA, which we found to be sufficient to induce chemoattractant production in macrophages. Localized synthesis of a chemoattractant in the aortic wall from a precursor released by PVAT could explain the observed PVAT-dependence of macrophage influx in the aorta (rather than to the PVAT itself) in response to CL treatment. Notably, we could find no evidence of increased inflammation in the aortas of CL-treated mice. Interestingly, we also found a significant increase in the n-3 fatty acid DHA in the supernatant of activated PVA, which potentially could promote and sustain the M2-like phenotype of the aortic macrophages via Resolvin D synthesis. This is particularly intriguing since an increase in both plasma AA and DHA has previously been shown to correlate with reduced aortic stiffness in humans ^38^. Importantly, 5-LOX is a rate-limiting enzyme in the synthesis of several PUFA-derived chemoattractants as well as pro-resolving lipid mediators, and is exclusively expressed by *Lyve1*^+^ aortic macrophages in the aorta. Interestingly, macrophages have previously been shown to rapidly become depleted of AA without continuous supply of exogenous polyunsaturated fatty acids ^39^, while adipocytes have recently been suggested to be a source of sustained AA production from linoleic acid ^40^. A local source of exogenous AA in close proximity to the aortic wall could therefore be important for sustained eicosanoid-based chemoattraction in the vascular wall during aortic remodeling. Importantly, simultaneous DHA-induced production of pro-resolving mediators could ensure that attracted monocytes arrive in a local milieu that promotes differentiation towards a remodeling/alternatively activated macrophage phenotype.

Exposure to cold temperatures has strong implications for central hemodynamics due to peripheral vasoconstriction leading to early wave reflection and increased augmentation index ^15^. This increases the systolic aortic blood pressure, potentially leading to increased hemodynamic stress on the aortic valves ^15^. Our data suggest a potential mechanism through which parallel activation of PVAT mitigates the detrimental cardiac effects of cold exposure on the peripheral vessels by improving the Windkessel effect of the ascending aorta, compensating for increased pressure augmentation and lowering the cyclic strain on the aortic valve leaflets.

The presence of active BAT in humans is associated with cardiometabolic health and reduced incidence of cardiovascular disease ^7^. Although the sample size in our cohort is limited by the low overlap of patients with detectable BAT activity, an AS ICD code, and available ^18^F-FDG PET-CT scans, we found a significant negative correlation of thermogenic mediastinal adipose tissue activity with diagnostic criteria of AS severity, such as peak jet velocity, in human patients. Mediastinal adipose tissue is defined as any adipose tissue found in the mediastinum, which may or may not be perivascular. However, mediastinal ^18^F-FDG uptake was recognized specifically as periaortic by radiologists as far back as 2005 ^41^, before metabolically active thermogenic adipose tissue was widely recognized to exist in adult humans ^42–44^. Since then, detailed ^18^F-FDG PET-CT studies have confirmed the metabolic activity of periaortic adipose tissue in humans ^45^. The periaortic adipose tissue at the human ascending aorta is small, and is unlikely to significantly contribute to whole-body temperature regulation ^46^.

However, our data indicate that PVAT could secrete factors that regulate remodeling and distensibility of the aortic wall. Interestingly, a recent study also showed that AS is associated with increased inflammation and reduced browning in periaortic fat, further highlighting the link between PVAT and AS progression ^46^.

While valvular wire injury is a state-of-the-art model for murine AS, it may not fully recapitulate the etiology of human AS, which is a gradual degenerative disease that takes many years to develop. However, it shares several key hallmarks with human AS, e.g. leaflet hypertrophy, immune cell infiltration, calcification, and fibrosis. Furthermore, it reliably induces echocardiographic findings mirroring those in human AS patients, such as increased peak jet velocity and transvalvular pressure gradient. Additionally, genetic ablation of PVAT by knockdown of *Pparγ* in smooth muscle cells may have unknown effects other on cells that co-express *Sm22α* and *Pparγ*. However, we corrected for this effect as much as possible by comparing the effects of e.g. CL316,243 administration or cold exposure in PVAT+ or PVAT- mice separately. As surgical removal of all PVAT surrounding major arteries is not feasible, the PVAT- knockout mouse model used in this study remains the best current model for delineating the role of PVAT in mice.

Our study shows that activation of PVAT ameliorates AS and is associated with an increased distensibility of the ascending aorta. Our results indicate that this is due to release of oxylipins and remodeling of the aortic wall. Importantly, we found that PVAT activity correlates with lower severity of AS in humans, suggesting a possible novel therapeutic avenue to treat AS by targeting PVAT.

## Author contributions

SH and AP designed the project and wrote the manuscript. KW performed mass spectrometry measurements. AS and ZS-R prepared samples for immunostaining of ascending aortas. JAB, NGB, and PC performed analysis of mediastinal BAT activity in AS patients. LRS supplied bone marrow-derived macrophages. ZS-R and SH performed single nucleus sequencing data from murine aortas. BF coordinated the import of the Sm22a^Cre^ mice, performed cold exposure experiments pertaining to the bulk mRNA sequencing of PVAT and aortic valves, and performed RNA extraction, mRNA sequencing, and partial sequencing analysis of these samples. SZ and GN provided infrastructure for the wire injury model. SH performed all other experiments. All authors contributed to reviewing and editing the final manuscript.

## Acknowledgements

The authors would like to thank Sandra Adler and Marta Stei for assistance with coordination of animal experiments; Gloria Martinac, Pia Fuchs, Christoph Schwelle and Henning Panatzek for technical assistance with the wire injury surgeries, and Patricia Zehner and Markus Raspe for technical assistance with histology. We thank Joachim Herz at UT Southwestern for providing the Sm22a-Cre mice ^14^ and Irmgard Förster for the PPARγ-floxed mice. We would also like to thank the Next Generation Sequencing Core Facilities of the Medical Faculty at the University of Bonn. Parts of the figures were created with BioRender.

## Data availability statement

Data supporting the findings of this study can be made available from the corresponding authors upon reasonable request.

