## Extended methods for "Activation of perivascular thermogenic fat improves aortic stenosis"

### **Bulk mRNA sequencing**

#### PVAT from cold-exposed mice:

PVAT samples from six mice were pooled for each replicate. RNA isolation was performed using InnuSOLV reagent. Library preparation and sequencing were performed by NGS Core Facility, Life & Brain Center, 53127 Bonn. For 3'mRNA-Seq, the library was prepared using QuantSeq 3' mRNA-Seq Library Prep Kit FWD for Illumina and sequencing was performed on HiSeq2500 V4 (High Output Mode, 1x50bp), following manufacturers' guidelines. The generated reads were single-end and of length of 50 nucleotides and were processed according to the Quant<sup>TM</sup>Seq User Guide. First 12 nucleotides, poly(A) tails, and nucleotides with Phred quality score below 30 were trimmed by Cutadapt. Additionally, minimum read length was set to 20 nucleotides. Next, the reads were aligned to a reference genome of a C57BL/6J mouse GRCm38.p6210 using STAR Aligner<sup>206</sup>. Afterwards, reads per gene were counted with HTSeq software. Until this point, each step was performed on Ubuntu 20.04 LTS operating system. The differential expression analysis was performed using DESeq2. GSEA analysis was performed with GSEA 4.3.3 (Broad Institute). Heatmaps were generated with pheatmap (https://github.com/raivokolde/pheatmap). For cold exposure, mice were housed at 18°C for 7 days for acclimation, followed by 4°C for 7 further days. Control mice were housed at 23°C.

#### Ascending aortas from wire-injured mice

RNA isolation was performed using InnuSOLV reagent. Library preparation, sequencing, and bioinformatic analysis was performed with Novogene GmbH (Munich, Germany):

#### *Library preparation for Transcriptome sequencing*

#### *Non strand specific library*

Messenger RNA was purified from total RNA using poly-T oligo-attached magnetic beads. After fragmentation, the first strand cDNA was synthesized using random hexamer primers followed by the second strand cDNA synthesis. The library was ready after end repair, A-tailing, adapter ligation, size selection, amplification, and purification. The library was checked with Qubit and real-time PCR for quantification and bioanalyzer for size distribution detection.

#### *Strand specific library*

Messenger RNA was purified from total RNA using poly-T oligo-attached magnetic beads. After fragmentation, the first strand cDNA was synthesized using random hexamer primers. Then the second strand cDNA was synthesized using dUTP, instead of dTTP. The directional library was ready after end repair, A-tailing, adapter ligation, size selection, USER enzyme digestion, amplification, and purification. The library was checked with Qubit and real-time PCR for quantification and bioanalyzer for size distribution detection.

#### *Clustering and sequencing*

After library quality control, different libraries were pooled based on the effective concentration and targeted data amount, then subjected to Illumina sequencing. The basic principle of sequencing is "Sequencing by Synthesis", where fluorescently labeled dNTPs, DNA polymerase, and adapter primers are added to the sequencing flow cell for amplification. As each sequencing cluster extends its complementary strand, the addition of each fluorescently labeled dNTP releases a corresponding fluorescence signal. The sequencer captures these fluorescence signals and converts them into sequencing peaks through computer software, thereby obtaining the sequence information of the target fragment.

#### *Bioinformatics Analysis Pipeline*

##### *Data quality control*

Raw data (raw reads) of fastq format were firstly processed through fastp software. In this step, clean data (clean reads) were obtained by removing reads containing adapter, reads containing poly-N and low quality reads from raw data. At the same time, Q20, Q30 and GC content the clean data were calculated. All the downstream analyses were based on the clean data with high quality.

##### *Reads mapping to the reference genome*

Reference genome and gene model annotation files were downloaded from genome website. Use HISAT2 (2.2.1) to build the index of the reference genome, and use HISAT2 to align paired-end clean reads to the reference genome. HISAT2 can use the gene model annotation file to create splice-aware alignments, providing better alignment accuracy compared to other non-splice alignment tools.

##### *Quantification of gene expression level*

featureCounts (2.0.6) was used to count the reads numbers mapped to each gene. And then FPKM of each gene was calculated based on the length of the gene and reads count mapped to this gene. FPKM, expected number of Fragments Per Kilobase of transcript sequence per Millions base pairs sequenced, considers the effect of sequencing depth and gene length for the reads count at the same time, and is currently the most commonly used method for estimating gene expression levels.

#### *Differential expression analysis*

For DESeq2 with biological replicates: Differential expression analysis for two conditions/groups was performed using the DESeq2 R package (1.42.0). DESeq2 provides statistical programs for determining differential expression in digital gene expression data using models based on negative binomial distribution. The resulting P-value is adjusted using the Benjamini and Hochberg's methods to control the error discovery rate. The threshold of significant differential expression : $\text{padj} \leq 0.05$ & $|\log_2(\text{foldchange})| \geq 1$. For edgeR without biological replicates: Prior to differential gene expression analysis, for each sequencing library, read counts were adjusted using the edgeR R package (4.0.16) by scaling normalization factors to eliminate differences in sequencing depth between samples, followed by differential expression analysis. The resulting P value is adjusted using the Benjamini and Hochberg's methods to control the error discovery rate. The threshold of significant differential expression: $\text{padj} \leq 0.05$ & $|\log_2(\text{foldchange})| \geq 1$.

#### *GO and KEGG enrichment analysis of differentially expressed genes*

Gene Ontology (GO) enrichment analysis of differentially expressed genes was implemented by the clusterProfiler (4.8.1), in which gene length bias was corrected. GO terms with corrected P-value less than 0.05 were considered significantly enriched by differential expressed genes. KEGG is a database resource for understanding high-level functions and utilities of the biological system, such as the cell, the organism and the ecosystem, from molecular-level information, especially large-scale molecular datasets generated by genome sequencing and other high-through put experimental technologies (http://www.genome.jp/kegg/). We used clusterProfiler R package to test the statistical enrichment of differential expression genes in KEGG pathways.

### High Performance Liquid Chromatography and mass spectrometry for eicosanoids and arachidonic acid

Cell culture media was thawed on ice. 200 µl of media was transferred to an Eppendorf tube and mixed with 100 µl of 0.5% (v/v) acetic acid, 50 µl of acetonitrile and vortexed thoroughly. 400 µl of Ethyl acetate/Hexane (20/80, (v/v)) was added and samples were centrifuged at 1550g for 3 minutes. The organic phase was transferred into a new Eppendorf tube and the solvents were evaporated in an Eppendorf speed vac at 30°C for 10 minutes. In total 600 µl of media per sample was extracted and pooled. Samples were reconstituted in 100% methanol and transferred into an LCMS glass vial and stored at 4°C for direct measurement. LCMS analysis was done with a Orbitrap Exploris120 mass spectrometer (ThermoFisher) coupled to a VanqishFlex HPLC (ThermoFisher). A Accucore UPLC C30, 2,6 µm, 2.1\*150 mm column (ThermoFisher) was used. For measurement of HETE-species and LTB<sub>4</sub>, the mobile phase was a 32 min gradient based on acetonitrile/water (50/50), with 5 mM ammonium formate + 0.1% formic acid (A) and Isopropanol/acetonitrile/water (85/10/5), with 5 mM ammonium formate + 0.1% formic acid (B) at the flow rate 0.25 ml/min. The gradient was as following: 10% B (0 min), 86% B (20 min), 95 % B (22 min), 95% B (26 min), 10% B (26.1 min), 10% B (32 min). For measurement of DHA derived species and untargeted analysis the mobile phase was a 26 min gradient based on water +0.1% formic acid (A) and acetonitrile/isopropanol (90/10) (B) at the flow rate 0.35 ml/min. The gradient was as following 20% B (0 min) to 35% B (2.5 min), 40 % B (4.5 min), 42 % B (6.0 min), 50 % B (8.0 min), 65 % B (14.0 min), 72.5 % B (15.5 min), 100 % B (16.8 min), 100 % B (17.8 min), 20 % B (18.0 min), 20% B (26 min). The mass spectrometer was operated in HESI negative mode with the following settings: spray voltage -3.5kV, sheath gas (40 a.u.), aux gas (5 a.u.), sweep gas (3 a.u.), ion transfer tube 230 °C, vaporizer temperature 350 °C with 120000 mass resolution for MS1 acquisition. DDA MS2 acquisition for compound identification was accomplished utilizing AquireX (ThermoFisher) with background ion exclusion and subsequent inclusion/exclusion lists with the following ddMS2 settings: isolation window (0.4 m/z), HCD normalized collision energy 30,60,80%, AGC target set to standard and automatic maximum ion injection time at 30000 mass resolution. Area integration and normalization via internal standardization for targeted compound analysis was done in TraceFinder 5.1. (ThermoFisher). Compound Discoverer SP 3.3 (ThermoFisher) was used for untargeted peak identification and area integration.

#### **Wire injury model**

Mice were anesthetized by intraperitoneal injection of ketamine (150 mg/kg) and xylazine (16 mg/kg), and the right carotid artery was exposed. Blood flow was stopped using ligature loops. A straight guide wire with a shortened and soldered tip (Abbott HI-TORQUE 0.014") was introduced into the left ventricle under echocardiographic guidance, passed over the aortic valve and advanced into the left ventricular apex and pulled back into the left ventricular outlet, just below the aortic valve level, and then rotated over the valve with a speed of two rotations per second. Intraperitoneal administration of buprenorphine (0.1 mg/kg) followed by buprenorphine supplementation in the drinking water was used as analgesia. In mice treated with CL316,243 (1 mg/kg/day), an osmotic minipump (Alzet 1004) was implanted three days after wire injury. Control mice were implanted with pumps containing NaCl. The analgesia regimen following pump implantation was intraperitoneal administration of buprenorphine (0.1 mg/kg) followed by supplementation of metamizole in the drinking water. For cold exposure, mice were acclimated at 16°C for three days followed by 7 days at 8°C prior to wire injury. After injury, the mice were allowed to recover for three days before returning to 8°C for the duration of the experiment. Mice with pre-existing aortic stenosis prior to wire injury (baseline peak velocity $\geq 2000$ mm/s) were excluded from analysis.

### **Histology**

#### Paraffin sections

Adipose tissues and aortas were fixed in PBS containing 4% PFA overnight and dehydrated using ethanol. The paraffin-embedded tissue was cut into 5 μM sections. Sections were blocked with 2% goat serum for 30 min at room temperature in tris-buffered saline. Immunohistochemistry (IHC) stainings were performed using a primary UCP1 antibody (1:75) (custom made, Thermo Fisher Scientific) overnight. Incubation with a secondary antibody conjugated with horseradish peroxidase (SignalStain Boost IHC; Cell Signaling, 8114S) was performed for 1 h at room temperature. Sections were visualized using 3,3'-diaminobenzidine (DAB) substrate (Vector Laboratories). For H&E staining, the sections were stained with haematoxylin (Merck, 1.09249) for 2 min and rapidly washed with distilled water. Staining with eosin (Merck, 1.09844) was performed immediately after by incubating the tissue section in eosin for 1 min and then washing for 4 min with distilled water.

#### Cryosections

Aortic valves and ascending aortas were fresh-frozen in OCT and post-fixed after sectioning using 4% PFA in PBS. H&E staining was performed as stated above. CD68 (origene SM1550PS, 1:100) staining was performed overnight followed by incubation with Alexa Fluor-conjugated secondary antibody (1:500) for 1 hour. Picrosirius red stain was performed using the Picro Sirius Red Stain Kit (Connective Tissue Stain, Abcam AB150681) according to the manufacturer's instructions.

### **Perivascular adipocytes**

Perivascular adipocyte precursor cells were isolated from thoracic PVAT of male C57BL/6J mice. PVAT was digested using Collagenase IV for 30 minutes with intermittent shaking. After centrifugation, the stromal vascular fraction was isolated and washed with MACS buffer. The washed cells were incubated with 10μl FcR blocking reagent (Miltenyi) for ten minutes at 4°C, and then incubated with 1μg of biotinylated anti-CD200 antibody (Thermo Fischer MA5-17979) for five minutes at room temperature. The labeled cells were then depleted using anti-biotin microbeads (Miltenyi 130-105-637) on LS columns according to the manufacturer's instructions. PDGFRα-positive cells were then isolated from the CD200-depleted fraction using CD140a microbeads (Miltenyi 130-101-502) according to the manufacturer's instructions. Eluted cells were seeded in 24-well plates and cultured in DMEM with 10% FCS, 10 ng/ml FGF-2, and 100 ng/ml Heparin sodium salt. After one day of culture, the cells were transduced with a lentiviral vector carrying the SV40 large T-antigen at a dose equivalent to 100 ng reverse transcriptase. After expansion and passaging, the cells were differentiated to mature adipocytes as previously described<sup>1</sup>.

### **Oxylipin secretion from perivascular adipocytes**

#### Sample preparation for LC-MS/MS

##### *Preparation of tissue samples and quality controls:*

100 μL of cell supernatant were transferred into 1.5 mL Eppendorf tube. QC pool samples were prepared in triplicates by taking out 100 μL from each study sample. The pool sample was subsequently mixed and 100 μL were transferred into 1.5 mL Eppendorf tubes. QC reference samples were prepared in triplicates in 1.5 mL

Eppendorf tubes by mixing 5 μL of the standard mixture (30 ng/mL for oxylipins and 300 ng/mL for PUFAs) with 45 μL of water. Blank (triplicate) and zero (single) samples were prepared by transferring 100 μL of cell culture medium into 1.5 mL Eppendorf tubes. Calibrators were prepared in 1.5 mL Eppendorf tubes by successive dilutions (factor 3) in water/MeOH (50:50, v/v) of the calibration mixture (200 ng/mL for oxylipins and 2000 ng/mL for PUFAs) to reach 9 calibrator points (cal.): 66.67 ng/mL (cal. 09) to 0.010 ng/mL (cal. 01). 30 μL of each cal. was then transferred to a new 1.5 mL Eppendorf tube.

Every tube was pre-cooled in wet ice before starting sample preparation and kept on wet ice all along the extraction procedure. For accurate quantification, 10 μL of 5 ng/mL (50 ng/mL for PUFAs) ISTD mixture were added to the samples, except zero sample.

##### *Extraction procedure:*

For lipid extraction, 150 μL of cold MeOH (-20 °C) were added to the samples followed by incubation for 10 min with vortexing every 3 min. Protein precipitation was performed by centrifugation of the samples at 10,000 x g for 15 min at 4 °C. The supernatant (around 150 μL) was transferred to a 1 mL Nunc<sup>TM</sup> 96-well polypropylene plate (ThermoFisher), and the volume was adjusted with water to reach 1 mL (final MeOH concentration of 15%) and mixed up. Solid phase extraction was then performed with a Strata-X Micro 96-well plate, 33 μm, 2 mL (Phenomenex) using a positive pressure-96 processor (Waters). After SPE plate conditioning with 2 x 0.5 mL MeOH and then 2 x 0.5 mL water, 2 x 0.5 mL of each sample were loaded on the SPE plate. After rinsing with 2 x 0.5 mL 10% MeOH in water (v/v) the analytes were eluted with 2 x 100 μL MeOH into a new 1 mL 96-well plate. Samples were transferred to a

select-a-vial 96-well plate with 300 μL glass inserts (Analytical Services) and evaporated to dryness at 30 °C with nitrogen gas. Analytes were resuspended with 30 μL 50% MeOH in water (v/v), vortexed, and centrifuged for some seconds at 1000 x g before direct injection into the analytical system.

##### *LC-MS/MS lipid mediator analysis*

All samples were measured with an Exion UHPLC-system coupled to a QTRAP 6500+ mass spectrometer (SCIEX, Darmstadt, Germany) operated with Analyst 1.6.3. Chromatographic separation was achieved using a Kinetex C18 reversed phase column (1.7 μm, 100 × 2.1 mm, Phenomenex) with a SecurityGuard Ultra Cartridge C18 (Phenomenex) precolumn, heated at 40 °C. Mobile phases A, water:ACN (70:30, v/v) + 100 μL AA, and B, ACN/IPA (50:50, v/v) were used with gradient program with an isocratic flow rate of 500 μL/min as follow: 0% B at 0 min, 70% B at 6.5 min, 100% B at 7.8 min, 100% B at 9.5 min, and 0% B at 11 min. The autosampler was operated at 4 °C with an injection volume of 10 μL of sample.

The coupled mass spectrometer was equipped with an electrospray ionization (ESI) Turbo-VTM source set to negative mode. Source parameters were optimized to the following values: source temperature 500 °C, curtain gas flow 40 psi, ionspray voltage –4000 V, ion source gas 1 50 psi, ion source gas 2 40 psi. Metabolites were analyzed via scheduled multiple reaction monitoring (sMRM) with nitrogen as collision gas. All MRM transitions were optimized for each compound, as well as the source parameters such as declustering potential, collision energy, cell exit potential and entrance potential. The sMRM detection window was set to 60 s. Acquisition time was about 8.5 min.

SciexOS software (Sciex) was finally used for peak detection, integration[MH1.1] and for quantitation of compounds (MQ4 algorithm). For quantification, linear calibration curves were generated from extracted calibrator samples for every compound via the IS method using the area ratio between the analyte and its ISTD, with a weighting factor of 1/x.

#### Data Processing

All data were processed using R 4.4.1.

A multistep procedure was used to ensure high data quality: First, lipids with a coefficient of variation larger than 25% in the QC-pool samples were discarded (n = 6). We then discarded lipids where missing values (NA) were observed in more than 50% per group in all investigated groups (n = 28).

The last quality control step comprised the calculation of the dispersion ratio (D-ratio) that uses the ratio of the technical variance (determined by QC-pool samples) and the total variance (variance of the biological samples) as a quality marker for each lipid <sup>2</sup>. We used a D-ratio cut-off of 50 %, as this implies that the technical variance is larger than the biological variance (n = 2 lipids were removed). After the quality control, n = 24 lipids remained in the data set.

Missing values (n = 82; equivalent to 34% of whole data set) were imputed using the GSimp imputation method approach as described by Wei et al. <sup>3</sup>. The algorithm was initialized using a quantile regression approach for left-censored missing data (QRILC). Missing values were then predicted using a Gibbs sampler approach using an elastic net model with alpha = 0.5 and lambda = 0.01. 50 iterations were used to optimize the value for each missing variable (iters\_each) and 10 iterations for the whole data set (iters\_all).

#### **Bone marrow-derived macrophages**

Bone marrow cells were isolated from the tibia and fibula of male C57BL/6J mice by centrifugation, and cultured in RPMI medium supplemented with 10% FCS, 1% L-Glutamine, and 15% m-CSF. After six days in culture, the cells were differentiated to alternatively activated macrophages by treatment with IL-4 (10 ng/ml) and IL-13 (10 ng/ml) for 24 hours. After differentiation, the macrophages were washed with Freestyle 293 medium and treated with 40 μM BSA-conjugated AA, DHA, or both, in Freestyle 293 medium for one hour, after which the supernatant was isolated and centrifuged at 1000 G for 10 minutes to remove cellular debris. Control cells were treated with an equivalent amount of BSA without AA/DHA.

#### **Single cell sequencing**

Ascending aortas from male C57Bl/6-J mice injected daily with either saline or 1mg/kg CL316,243 for 10 days were dissected and snap-frozen in liquid nitrogen. Aortic segments from 5 mice were pooled for each condition. Frozen tissues were thawed, lysed in CHAPS buffer, and homogenized in a Dounce homogenizer. Lysates were filtered through a 40 μm sieve, and nuclei were collected by centrifugation for 5 minutes at 500 G. Nuclei were resuspended to a concentration of 1000 nuclei per μL. Library preparation was then conducted using the 10X platform (10X Genomics) according to the manufacturer's instructions. Sequencing was performed by a commercial provider (Novogene GmbH (Munich, Germany)). Data was processed using the Seurat package <sup>4</sup>.

#### **Statistical analysis**

Data are represented as means ± SEM. Unless otherwise stated, comparisons of single variables across more than two conditions were performed using one-way

ANOVA and Tukey's post-hoc test. Comparisons of more than one variable across more than two conditions were performed using two-way ANOVA and Sidak's post-hoc test. Comparisons between two conditions were performed using two-tailed Student's t-test. Outliers were excluded using ROUT with Q=1%. Statistical analysis was performed in GraphPad Prism (GraphPad Software, Inc, La Jolla, CA).

1. Haas B, Mayer P, Jennissen K, Scholz D, Berriel Diaz M, Bloch W, et al. Protein kinase g controls brown fat cell differentiation and mitochondrial biogenesis. *Sci Signal*. 2009;2:ra78
2. Broadhurst D, Goodacre R, Reinke SN, Kuligowski J, Wilson ID, Lewis MR, et al. Guidelines and considerations for the use of system suitability and quality control samples in mass spectrometry assays applied in untargeted clinical metabolomic studies. *Metabolomics*. 2018;14:72
3. Wei R, Wang J, Jia E, Chen T, Ni Y, Jia W. Gsimp: A gibbs sampler based left-censored missing value imputation approach for metabolomics studies. *PLoS Comput Biol*. 2018;14:e1005973
4. Satija R, Farrell JA, Gennert D, Schier AF, Regev A. Spatial reconstruction of single-cell gene expression data. *Nat Biotechnol*. 2015;33:495-502
