## Supplemental figures for "Activation of perivascular thermogenic fat improves aortic stenosis"

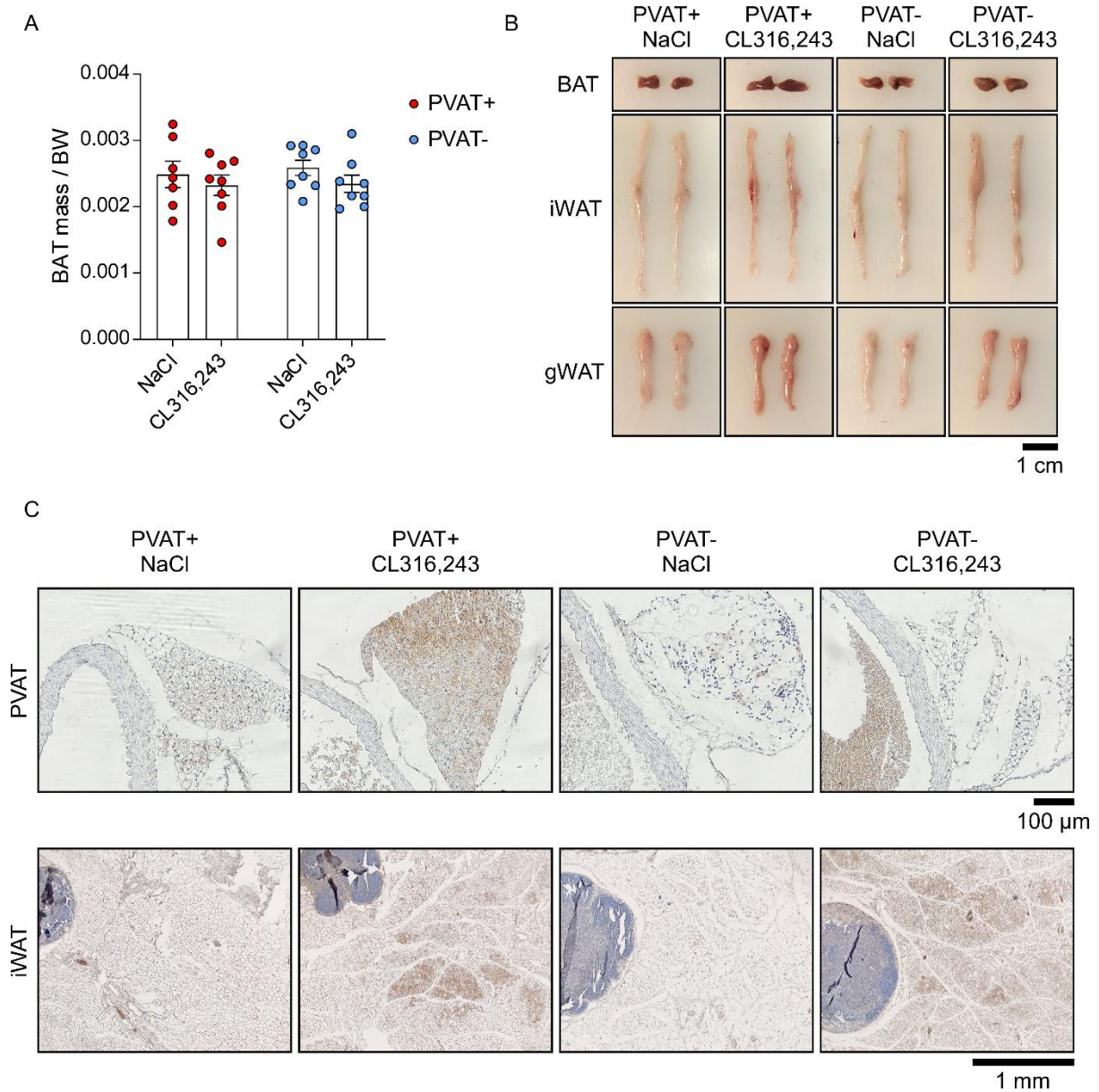

**Supplemental Figure 1: BAT mass and browning in PVAT+ and PVAT- mice. A:** BAT mass per body weight in NaCl- or CL-treated PVAT+ and PVAT- mice. $n=7-8$. **B:** Photographs of adipose tissues from NaCl- or CL-treated PVAT+ and PVAT- mice. Scale bar: 1 cm. **C:** UCP1 staining (brown) of PVAT and iWAT of NaCl- or CL-treated PVAT+ and PVAT- mice. Scale bar PVAT: 100 μm. Scale bar iWAT: 1 mm.

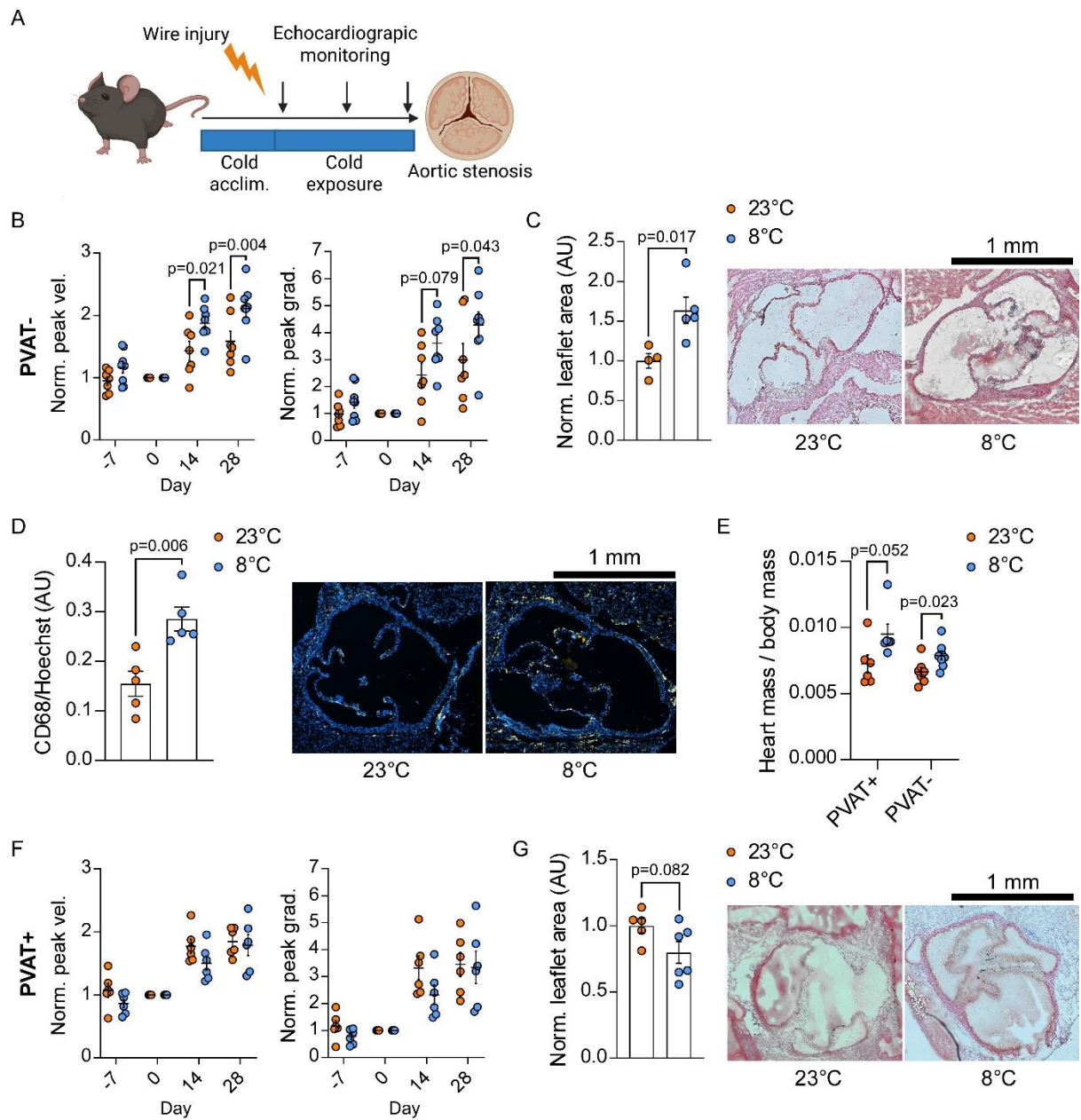

**Supplemental Figure 2: Effect of cold-induced PVAT activation on AS in PVAT+ and PVAT- mice.** **A:** Schematic illustrating the experimental layout. **B:** Peak jet velocity and peak gradient in $PPAR\gamma^{fl/fl}$ $Sm22a^{Cre/+}$ (PVAT-) mice at different timepoints of the experiment. $n=7-8$. Two-way ANOVA with Sidak's post-hoc test. **C:** Histological (H&E) analysis of leaflet area in wire-injured PVAT- mice. $n=4-5$. $t$-test. Scale bar: 1 mm. **D:** Immunohistological analysis of CD68-expressing cells in the aortic valve leaflets of PVAT- mice following wire injury. Blue: Hoechst, orange: CD68. $n=5$. $t$-test. Scale bar: 1 mm. **E:** Heart mass to body mass ratio of PVAT+ and PVAT- mice. $n=6-8$. $t$-test. **F:** Peak jet velocity and peak gradient in $PPAR\gamma^{fl/fl}$ $Sm22a^{+/+}$ (PVAT+) mice at different timepoints of the experiment. $n=6$. Two-way ANOVA with Sidak's post-hoc test. **G:** Histological (H&E) analysis of leaflet area in wire-injured PVAT+ mice. $n=5-6$. $t$-test. Scale bar: 1 mm.

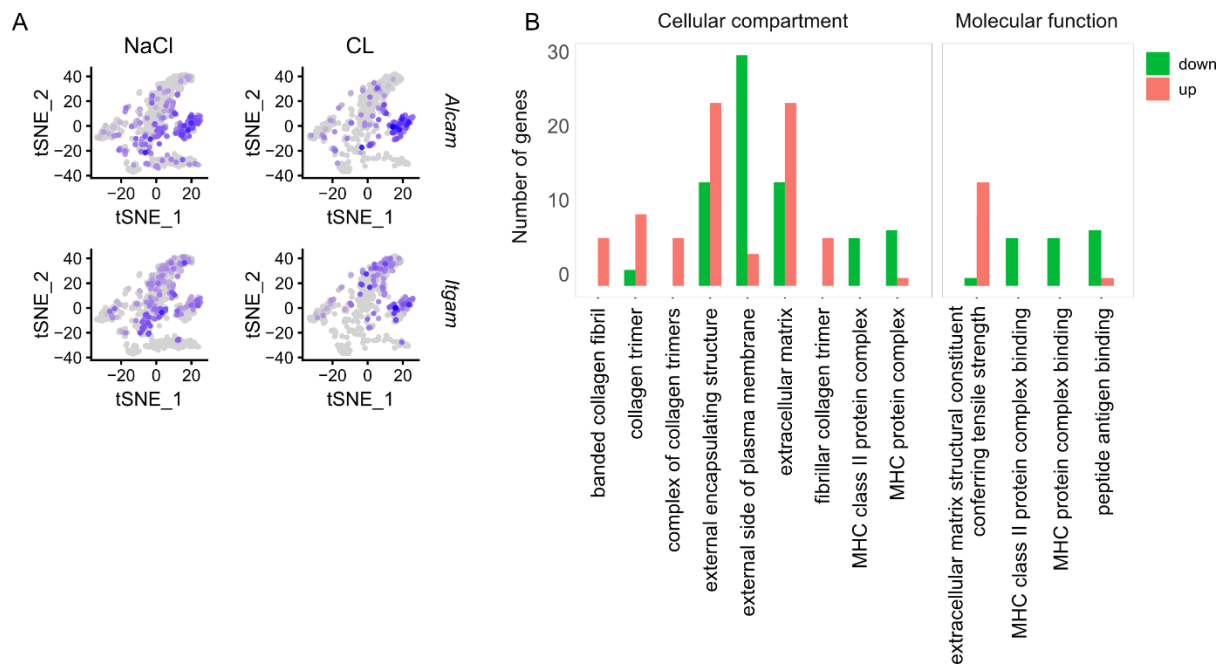

**Supplemental figure 3: A:** Feature plots showing expression of Alcam and Itgam in immune cell clusters of aortas from mice treated with either NaCl or CL316,243. **B:** Gene ontology analysis comparing genes regulated by CL treatment in the aortic wall of PVAT+ and PVAT- mice.

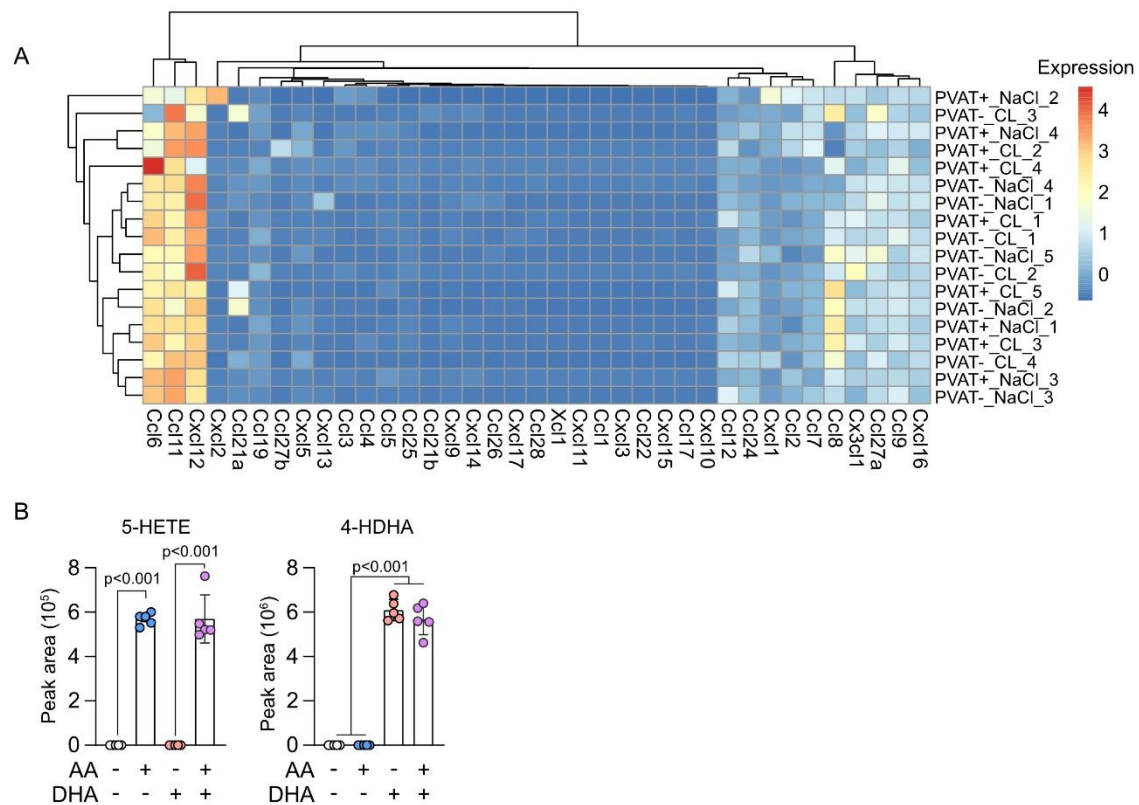

**Supplemental Figure 4. A:** Heatmap showing unsupervised clustering of chemokine expression in the ascending aortas of PVAT+ and PVAT- mice treated with either NaCl or CL. $n=4-5$. **B:** 5-HETE and 4-HDHA in the supernatants of BMDM treated with either AA or DHA or both. $n=5$. One-way ANOVA with Tukey's post-hoc test.
